# Dynamic interactions between top-down expectations and conscious awareness

**DOI:** 10.1101/151019

**Authors:** Erik L. Meijs, Heleen A. Slagter, Floris P. de Lange, Simon van Gaal

**Author notes:** Corresponding author: Simon van Gaal, University of Amsterdam, Department of Psychology, 1001 NK Amsterdam, the Netherlands.

## Abstract

It is well known that top-down expectations affect perceptual processes. Yet, remarkably little is known about the relationship between expectations and conscious awareness We address three crucial questions that are outstanding: 1) How do predictions affect the likelihood of conscious stimulus perception?; 2) Does the brain register violations of predictions nonconsciously?; and 3) Do predictions need to be conscious to influence perceptual decisions? We performed three experiments in which we manipulated stimulus predictability within the attentional blink paradigm, while combining visual psychophysics with electrophysiological recordings. We found that valid stimulus expectations increase the likelihood of conscious access of stimuli. Furthermore, our findings suggest a clear dissociation in the interaction between expectations and consciousness: conscious awareness seems crucial for the implementation of top-down predictions, but not for the bottom-up generation of stimulus-evoked prediction errors. These results constrain and update influential theories about the role of consciousness in the predictive brain.

---

A rapidly growing body of work indicates that sensory processing is strongly influenced by the expectations that we have about likely states of the world. Such expectations are shaped by the immediate environment or context in which we are operating, but also by learning, past experience and our genetic makeup ^1–3^. Expectations are typically thought to originate from higher-level brain regions, such as the prefrontal cortex, which may guide information processing in lower-level sensory regions via top-down (descending) projections ^4,5^. In this framework, what we consciously see is proposed to be strongly influenced by the brain’s expectations about, or its best guess of, the outside world ^6,7^. Initial studies support a tight relationship between expectations and conscious perception. For example, it has been shown that objects that are unexpected in a particular visual scene (e.g. a hammer in the kitchen) are detected more slowly than expected objects (e.g. a knife in the kitchen) ^8^. Further, a correct prediction about the nature of an upcoming stimulus (e.g. the orientation of a Gabor) improves its discrimination ^9^ and sharpens its neural representation ^10^. The brain can thus use predictive information in the environment to build expectations of stimulus frequency or conditional probabilities to modify subsequent sensory information processing and perception. These ideas have been formalized in several theoretical models, such as predictive coding and sequential sampling models ^3,11,12^. Although these frameworks are attractive in their simplicity and are rapidly growing in scientific stature, how exactly predictions shape conscious perception, and to what extent awareness guides prediction formation, is still speculative and largely unknown.

At present, there are several issues that need to be resolved in order to further our understanding of the relationship between predictions and consciousness. In this report we address three crucial issues. The first issue relates to the effect that predictions may have on conscious awareness itself. Regarding this, there are two opposing hypotheses. One possibility is that predictable inputs are cancelled from perception ^3,13^, leading to prioritized processing of unexpected input, reflected in an increased “prediction error” ^10,14–16^. As conscious perception may be directly linked to the amount of neural activity that a stimulus evokes ^17,18^, subjects may be more aware of unpredicted compared to predicted stimuli ^19^. Indeed, a study by Mudrik and colleagues ^20^ showed that stimuli that are not predicted by their spatial context (e.g. a checkerboard in the oven) break through continuous flash suppression faster than congruent stimuli (e.g. a cake in the oven). Alternatively, because valid predictions lead to minimization of prediction errors and improved stimulus representations ^10,21^, a diametrically opposed hypothesis states that conscious access may be boosted for input that is predicted. Support for this notion is provided by studies that have shown that valid predictions increase the speed of conscious access ^22–26^ and may help selecting or facilitating stimulus interpretation when (visual) input is ambiguous or noisy (Aru, Rutiku, Wibral, Singer, & Melloni, 2016; Bar et al., 2006; Chang, Kanai, & Seth, 2015; Panichello et al., 2013). It is yet an open question whether predictions can boost an otherwise unseen stimulus into conscious awareness, thereby enabling the switch from a nonconscious to a conscious stimulus representation, instead of just facilitating its cognitive interpretation or its speed of appearance in time.

The second question is to what extent prediction errors can be registered outside of conscious awareness. There is some evidence that simple or low-level prediction errors can be detected nonconsciously. For instance, it has been shown that “oddball” stimuli, which are often simple violations in auditory tone sequences (AAAAA vs. AAAAB), elicit early mismatch responses in electrophysiological signals, called the mismatch negativity (MMN) ^29,30^. Interestingly, such an MMN can even be observed when subjects are attentionally distracted from the violations in tone sequences by a demanding visual task ^31^. Further, even in several reduced states of consciousness, such as sleep ^32,33^, anesthesia ^34^ and vegetative state ^31,35^, an MMN can be observed. Therefore, it has been hypothesized that the MMN may reflect a low-level pre-attentive prediction error signal that is generated nonconsciously ^36–38^. However, it remains debated whether these mismatch effects actually reflect nonconscious prediction detection or whether these neural signals merely reflect a passive low-level sensory adaptation mechanism ^39,40^. This is mainly because the majority of the studies employing oddball-like paradigms in reduced states of consciousness have not dissociated prediction from adaptation, because deviant stimuli occurred both rarely in general (reflecting adaptation) and had a low probability based on the just preceding stimulus (reflecting prediction). The one study in which the authors aimed to dissociate these two mechanisms in an nonconscious state showed that adaptation mechanisms remain operative during sleep, whereas the prediction errors disappear ^32^. These latter results suggest that prediction errors may not be registered outside of conscious awareness. However, in reduced states of consciousness, control over the content of perception is strongly related to arousal-related mechanisms that are affected at the same time. It is therefore unknown whether prediction errors can be observed when the content of consciousness is exclusively manipulated ^41^ without affecting these mechanisms.

The final issue concerns the role of conscious awareness in *implementing* predictions. An assumption of many prediction-based models is that predictions are implemented via top-down (descending) neural activation. Interestingly, influential theories of consciousness suggest that conscious access requires similar top-down interactions between higher-level (e.g. prefrontal) and lower-level (e.g. visual) brain regions, referred to as feedback or recurrent processing ^42–44^. Information that does not reach conscious access is thought to only trigger feedforward activity or local recurrent interactions between posterior brain regions. As a result, it is unclear how nonconscious information, in the absence of feedback signals from prefrontal cortex, could lead to the implementation of predictions. At present, it thus remains an open question whether conscious awareness is needed in order for predictive information to influence perceptual processes, or whether predictions can be implemented outside of awareness.

In this report, we address the three above outlined questions in a set of three studies employing a combination of behavioral psychophysics and electroencephalographic recordings (EEG).

## Results

### Experiment 1: (how) do predictions affect conscious access?

In the first experiment we addressed the question if predictions about the likelihood of stimulus identity modulate the likelihood of conscious access, and if so, in what direction. To do so, we used the attentional blink paradigm ^45^. The attentional blink is an impairment in the conscious perception of the second of two target stimuli that are presented in rapid succession (RSVP: rapid serial visual presentation). Here we modified the paradigm in such a way that the first target (T1: the letter G or H, in green) predicted which of the second targets would most likely be presented (T2: the letter D or K, predicted=60%, unpredicted=20%, in white, Fig. 1A). On 20% of trials we presented a random distractor letter instead of a T2 target. At the end of each stream of letters, participants gave three responses. First, they indicated whether or not they had seen any of the two T2 targets (“seen”/”unseen” response). Second, they were prompted to make a forced-choice judgment about the identity of T2 (whether the letter D or K was presented). Third, participants had to make a similar forced-choice decision about the identity of T1 (whether the letter G or H was presented) (see Methods for details). Participants were not explicitly instructed about the predictive relationship between T1 and T2.

**Figure 1.**
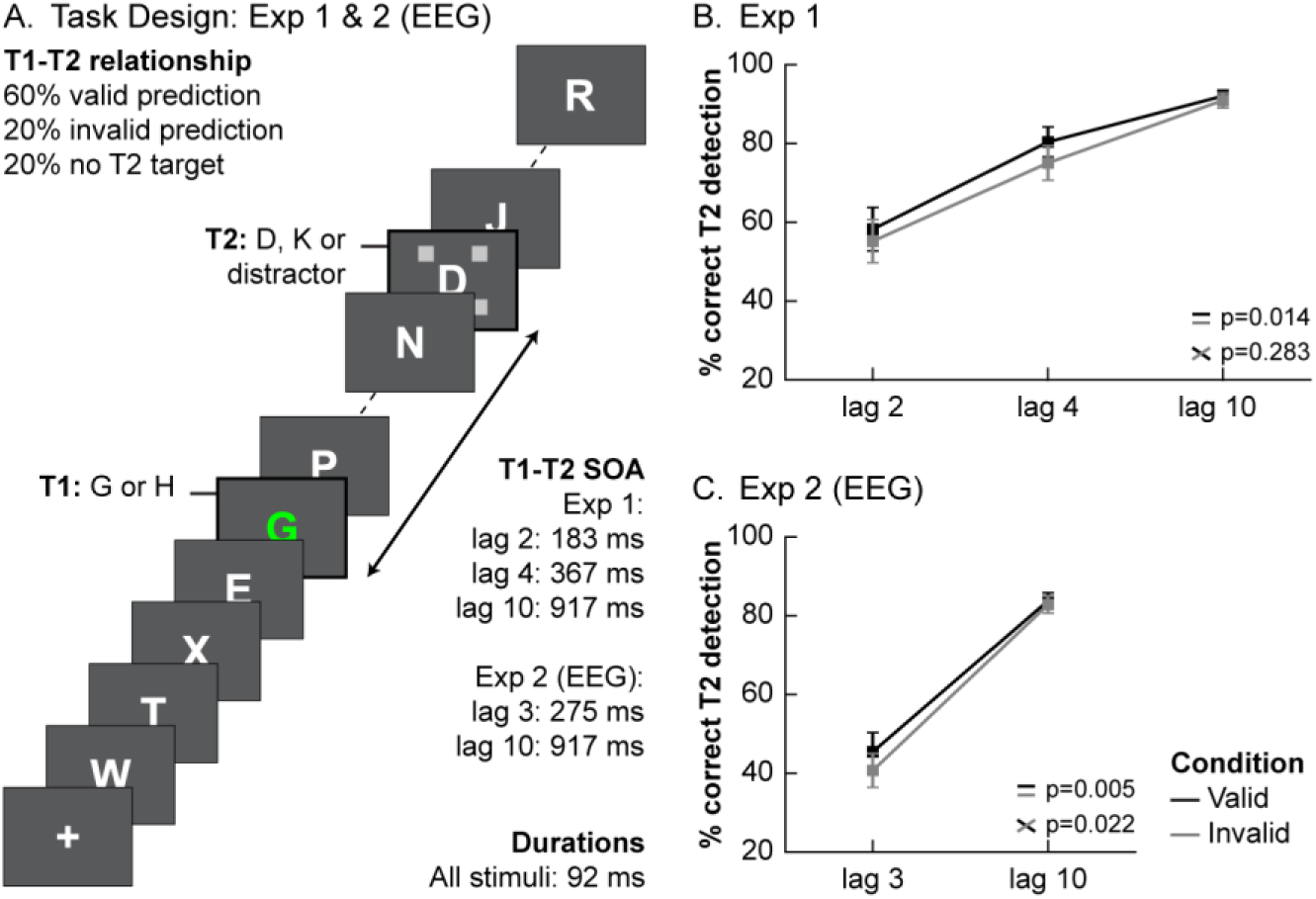
Task design and behavioral results of Experiment 1 and 2. **(A)** The trial structure of the attentional blink task used in Experiment 1 and 2. Each trial consisted of a stream of rapidly presented letters in which predefined target letters had to be detected and then reported at the end of the stream. The first target (T1: a green G or H) always appeared at the fifth position. The second target (T2: D or K), was presented at varying SOAs (lags) after the first one and was marked by placeholders. The identity of T1 predicted which of the T2 targets was most likely to appear, thereby introducing validly and invalidly predicted T2 targets. On 20% of the trials no second target was presented and a random distractor letter was presented instead. **(B)** Percentage of T2 target detection at each of the T1-T2 lags, after a valid or invalid prediction for Experiment 1. Validly predicted T2’s were significantly more often perceived than invalidly predicted T2’s. **(C)** Percentage of T2 target detection at each of the T1-T2 lags, after a valid or invalid prediction for Experiment 2. Again, validly predicted T2’s were more often perceived, in particular at short lags. Error bars represent SEM.

In Figure 1 we plot the percentage of trials in which T2 was correctly detected and T1 discrimination was also correct (average T1 accuracy was 94.20%, sd=5.77%) for the three different lags (lag 2, 4 and 10). T2 was considered to be detected correctly when participants indicated to have seen it (based on the first response) and correctly identified it (based on the second response). Overall, there was a clear attentional blink, as reflected by reduced T2 detection when the time (i.e. lag) between T1 and T2 was shorter (Figure 1B, main effect of lag: F_2,48_=48.15, p<0.001). Importantly, predictions modulated T2 detection rate. T2 detection was significantly better when T1 validly predicted T2 (black lines) compared to when the prediction was invalid (gray lines, main effect of validity: F_1,24_=7.10, p=0.014, no significant interaction between lag and validity: F_2,48_=1.30, p=0.283). These results extend several previous studies (Chang et al., 2015; Melloni et al., 2011; Pinto et al., 2015; Stein et al., 2015; Stein & Peelen, 2015) by showing that conscious perception is (partly) determined, in an “all-or-none” fashion, by the transitional probability of the input the brain receives.

While these data support the notion that valid expectations trigger access to consciousness, it has been recognized that such findings may not solely be due to changes in perception, but perhaps (also) due to changes in decision criteria or response biases ^46–48^. To rule out the possibility that our effects could be explained by a response bias in which people simply report the target letter that they expected based on T1, irrespective of whether they consciously perceived T2, we performed an analysis with T2 awareness (instead of T2 discrimination, see Methods) as the dependent variable. This analysis takes into account only participants’ first response (the “seen”/”unseen” response), regardless of whether subsequent T2 letter identification was correct or not. Crucially, this analysis cannot be influenced by any decision/response biases because the response was orthogonal to the participants’ expectation, and we still observed a qualitatively similar pattern of results (main effect of validity: F_1,24_=5.47, p=0.028). This finding highlights that validity truly boosted participants conscious access of T2, instead of merely eliciting a shift in response bias.

### Experiment 2: EEG markers of conscious and nonconscious prediction violations

Subsequently, we tested whether prediction violations can be elicited by nonconsciously processed unpredicted stimuli or whether conscious perception of a stimulus is a prerequisite for it to trigger neural prediction error responses. To test this, we measured subjects’ brain activity with EEG while they performed a similar task as in Experiment 1. First, we replicated the behavioral effects of Experiment 1 (Fig. 1C). Overall, T1 performance was high (M=93.61%, sd=7.31%) and T2 detection was higher at lag 10 than at lag 3 (main effect of lag: F_1,28_=128.72, p<0.001), reflecting a robust attentional blink. More importantly, validly predicted T2s were discriminated better than invalidly predicted T2s (main effect of validity: F_1,28_=9.49, p=0.005). In this experiment, the validity effect was significantly modulated by lag (validity x lag: F_1,28_=5.86, p=0.022), an effect that was numerically similar, but not significant in Experiment 1. Participants performed better for valid than invalid trials at lag 3, but there was no convincing evidence for an effect of predictions at lag 10 (lag 3 validity effect: t_28_=3.40, p=0.002; lag 10 validity effect: t_28_=0.98, p=0.334). Thus, effects of predictions were larger in the time window in which T2 more often goes unperceived.

Next, we investigated potential differences in the neural processing of predicted and unpredicted stimuli, as a function of stimulus awareness. To this end, we contrasted invalidly and validly predicted T2s and tested this difference using cluster-based permutation testing, correcting for multiple comparisons across both time (0- 750 ms) and (electrode) space (see Fig. 2 and Methods) ^49^. We found one significant difference over frontocentral electrode channels, which reflected greater T2-elicited negativity for invalid compared to valid trials between 174-314 ms (p=0.015, Fig. 2B), therefore potentially reflecting a type of mismatch response. We then further analyzed this component to test whether this difference was modulated by, or dependent on, conscious perception of T2. Crucially, the size of this fronto-central mismatch component was independent of T2 awareness (F_1,28_=0.04, p=.850, BF=0.254, Fig. 2C), indicating that both seen and unseen T2’s generated a fronto-central mismatch response.

**Figure 2.**
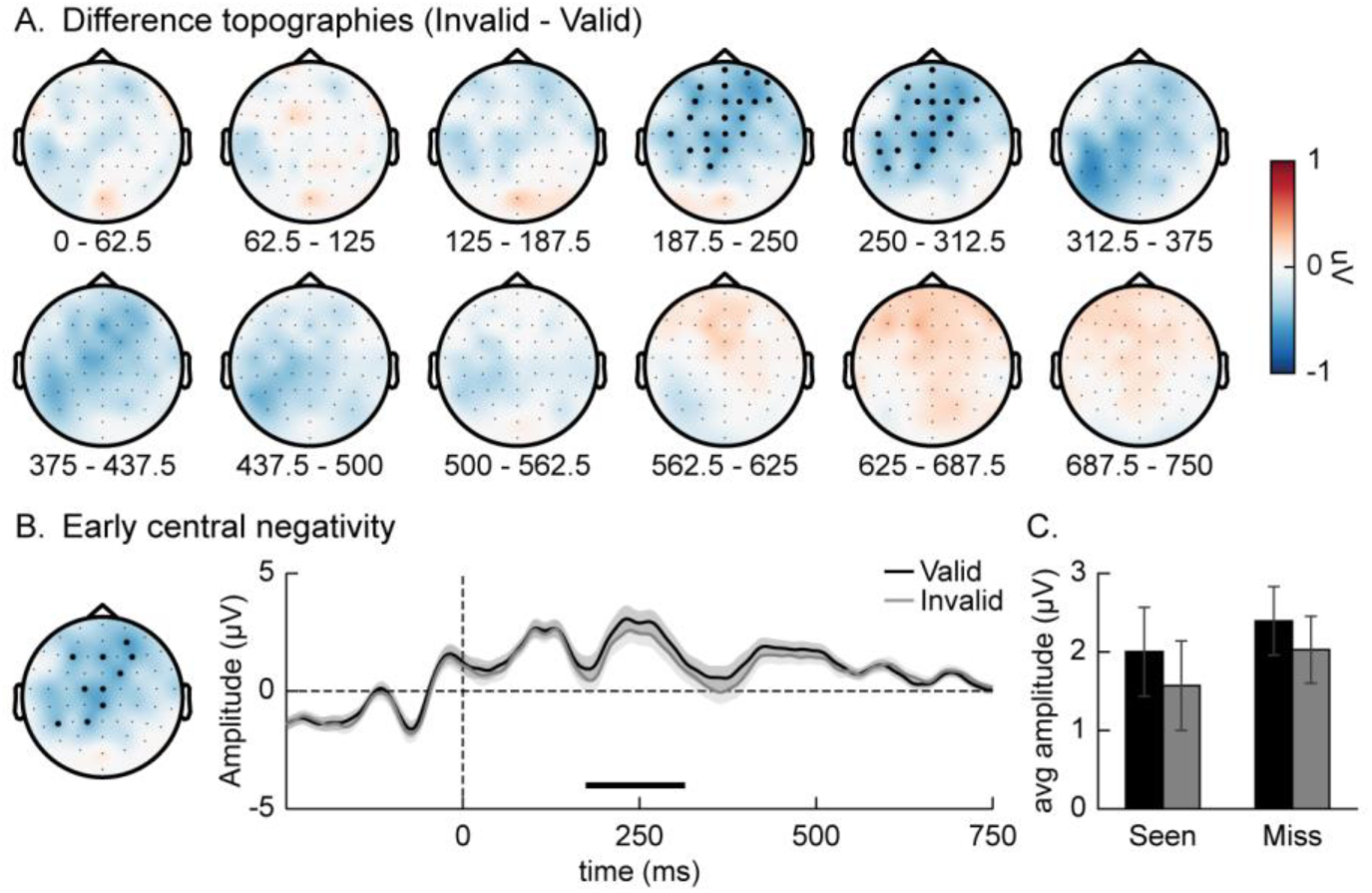
ERP effects. **(A)** Topographic maps of the difference between validly and invalidly predicted T2s over time (0 = T2 onset). Cluster-based permutation tests were used to isolate the significant events, while correcting for multiple comparisons across time and (electrode) space. On each head map, channels with a significant effect for at least 50% of its time window are highlighted. **(B)** The average ERP time-course of the 10 channels shown on the headmap on the left, shown seperately for each validity condition. The significant time-window is marked by a black line above the x-axis. Invalidly predicted T2s were associated with greater fronto-central negativity than validly predicted T2s. **(C)** Bar graphs showing the average amplitude of the four conditions (visibility x prediciton) for the significant neural event shown in B. In all plots error bars represent SEM.

Additionally, we also replicated previously reported findings related to T2 visibility ^50–52^, which we have detailed in the Supplementary Results section. In summary, seen versus unseen targets were associated with an increase in a posterior negative component that was later followed by a larger frontal and central positive component, reflecting larger N2/N3, P3a and P3b ERP components to seen targets. Within none of those components did we find that the T2 visibility effect was dependent on the validity of the prediction (all p>0.15). Finally, we directly tested for an interaction between conscious access and prediction by comparing the validity effect for T2 seen and T2 missed trials in a cluster-based permutation test. Again, no significant interactions between were found (all clusters p>0.10).

### Experiment 3: the role of conscious awareness in implementing top-down predictions

In our final experiment, we addressed the question whether prediction formation itself can unfold in the absence of awareness and subsequently influence conscious access (Fig. 3). To address this question, we changed the color of T1 from green to white and for each subject staircased T1 duration in such a way that T1 was correctly identified on approximately 75% of the trials (actual T1 identification performance: M=76.03%; sd=8.65%). T1 duration did not differ between trials where T2 was seen and trials where T2 was missed (T2 detection: t66=0.31, p=0.752; T2 seen: M=117.42 ms; T2 missed: M=117.46 ms), which indicates that T1 visibility was mainly determined by fluctuations in T1 processing, and not stimulus duration per se. Moreover, on 20% of trials no T1 was presented (but replaced by a distractor). Further, to test to what extent explicit knowledge of the predictive relationships between stimuli would increase the validity effects, we varied the moment at which explicit information about the predictive relations between T1 and T2 was provided. The experiment consisted of a training session and a test session on separate days. A first group of subjects received no explicit instructions about the predictive relations in either session and had to learn them implicitly through experience with the task; the second group received explicit instructions about the T1-T2 relations in the test session only, but not in the first training session; and the third group received explicit instructions already from the start of the experiment.

**Figure 3.**
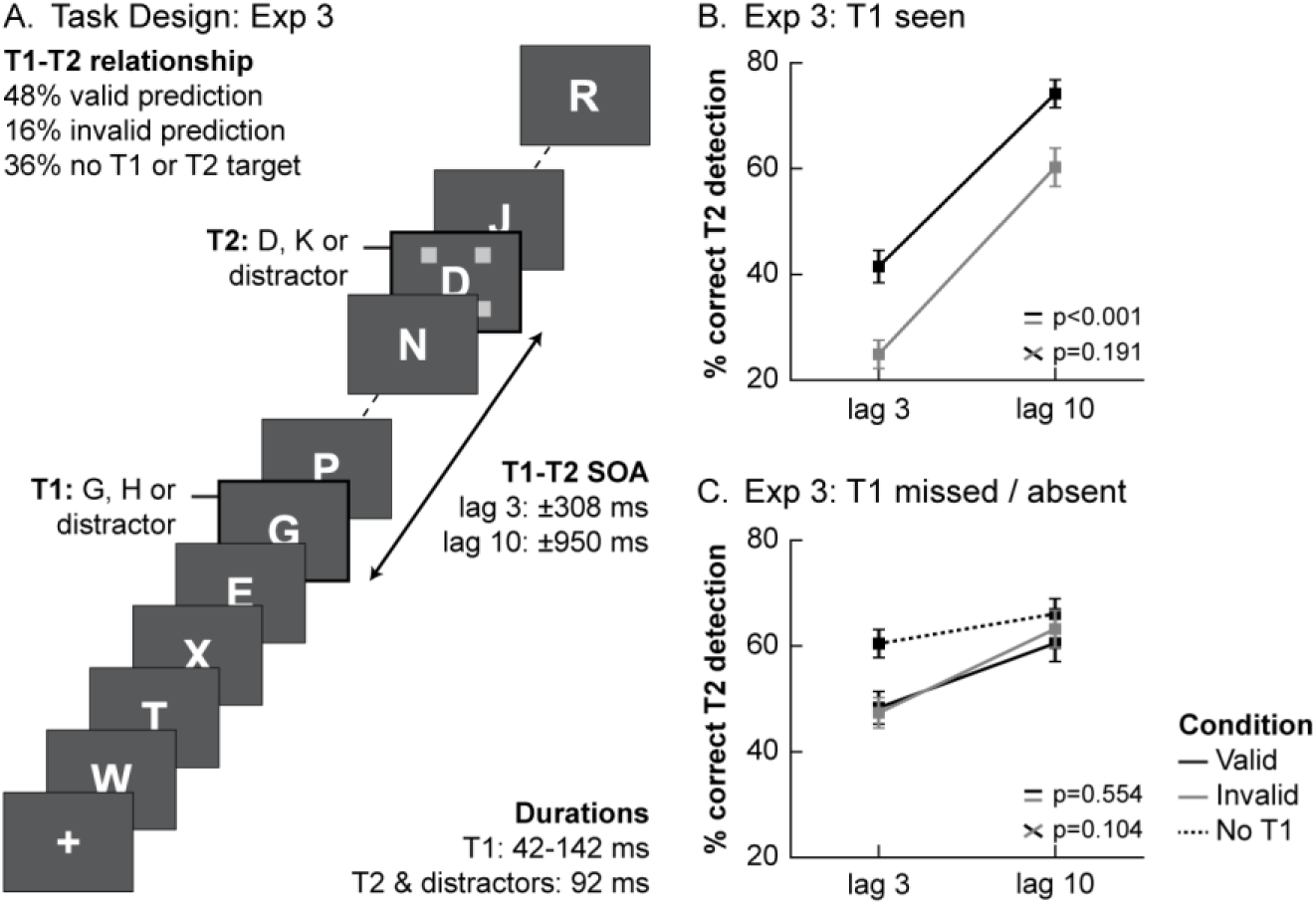
Task design and behavioral results of Experiment 3. **(A)** Trial structure of the task used in Experiment 3. T1 visibility was staircased at approximately 75% correct by manipulating its duration (on 20% of trials no T1 was presented). **(B)** Percentage of T2 target detection at each of the T1-T2 lags, after a valid or invalid prediction, for trials where T1 was correctly reported (T1 seen). As in Experiment 1 and 2, when T1 was seen, validly predicted T2’s were more often detected than invalidly predicted T2’s. **(C)** Solid lines show percentage of T2 target detection at each of the T1-T2 lags, after a valid or invalid prediction, for trials where T1 was presented but missed. In contrast to T1 seen trials (B), when T1 was not seen, validity did not enhance T2 detection. However, a missed T1 still triggered a significant attentional blink, as compared to trials on which no T1 was presented (dotted line). Error bars represent SEM.

T1 visibility strongly affected T2 detection. When T1 was seen, T2 detection was markedly lower than when T1 was missed (main effect of T1 awareness: F_1,64_=4.62, p=0.035), in particular at short lags (T1 awareness x lag: F_1,64_=72.95, p<0.001). Validly predicted targets were detected more often (main effect of validity: F_1,64_=33.39, p<0.001). The effect of prediction validity on T2 detection varied as a function of T1 awareness and instructions (T1 awareness x validity: F_1,64_=40.55, p<0.001; validity x instruction: F_1,64_=5.91, p=0.004; T1 awareness x validity x instruction: F_2,64_=11.33, p<0.001). When T1 was seen (Fig. 3B), a clear attentional blink was observed (main effect of lag: F_1,64_=170.01, p<0.001) and validly predicted targets were more often detected than invalidly predicted targets (main effect of validity: F_1,64_=64.97, p<0.001) (as in Experiment 1 and 2). Interestingly, we also observed a significant attentional blink for missed T1’s, reflecting a nonconsciously elicited AB (main effect of lag: F_1,64_=74.42, p<0.001). This AB effect cannot be explained by an overall T2 detection performance benefit for targets that are presented later in the trial because the AB was larger for trials on which T1 was presented but missed compared to trials on which no T1 was presented in the trial (lag x T1 presence: F_1,66_=24.19, p<0.001). However, although missed T1’s triggered an AB, prediction validity did not affect T2 detection performance for missed T1’s (main effect of validity: F_1,64_=0.35, p=0.554), regardless of the type of instruction participants received about the predictive relation between T1 and T2 (validity x instruction: F_2,64_=0.64, p=0.533). A Bayesian equivalent of the repeated-measures analysis strongly suggested validity should not be included in a model of the data (BF=.084).

The above results highlight that only when T1 was seen, valid predictions facilitated T2 detection. A post-hoc analysis on T1-seen trials only revealed that this effect was modulated by how explicitly we instructed participants about the predictive relationship between T1 and T2 (validity x instruction: F_2,64_=14.83, p<0.001). The validity effect, as defined by the difference between valid and invalid trials, averaged across the two lags, increased with more explicit instructions (group 1: 1.87%, group 2: 19.53%, group 3: 26.27%). These results indicate that, not only does the visibility of T1 define the predictive impact on T2 detection, but also the extent to which these predictive relations are (explicitly) known affects the impact of predictions on conscious access. This may also explain why the validity effect appeared more pronounced in Experiment 3 compared to Experiments 1 and 2, because in experiment 1 an d 2 subjects were not explicitly instructed about the predictive relations between T1 and T2.

Finally, in contrast to Experiment 2, on T1 seen trials the validity effect was independent of lag (validity x lag: F_1,64_=1.750, p=0.191). Since we anticipated stronger expectation effects at short lags, behavioral data from all three experiments was combined in a post-hoc analysis. Only trials on which T1 was correctly identified were used and for Experiment 1 we averaged data for lag 2 and lag 4 to create an average “short lag” condition. A significant interaction between validity and lag showed that across all experiments, the prediction effect was stronger at short lags compared to the long lags (validity x lag: F_1,118_=5.73, p=0.018; no validity x lag x experiment interaction: F_2,118_=0.065, p=0.937).

## Discussion

In this report we investigated three important questions regarding the intricate relationship between top-down expectations and conscious awareness. The first question that we addressed was how prior information about the identity of an upcoming stimulus may influence the likelihood of that stimulus entering conscious awareness. Using a novel attentional blink paradigm in which the identity of T1 cued the likelihood of the identity of T2, we showed that, stimuli that confirm our prediction have a higher likelihood of gaining access to conscious awareness than stimuli that violate our predictions, especially at short lags. This effect could not be explained by simple shifts in response criterion, suggesting that valid predictions amplify the perceptual strength of a stimulus and therefore increase the chance of conscious access, possibly due to the sharpening of its neural representations ^10^. This interpretation is supported by previous experiments that have shown that prior knowledge increases the speed (Chang et al., 2015; De Loof et al., 2016; Melloni et al., 2011; Pinto et al., 2015) and accuracy ^23^ of stimulus detection, extending these previous reports by showing that prior knowledge accelerates conscious access. Furthermore, they complement recent studies showing that the AB can be reduced when there is knowledge about temporal statistics of the task ^53,54^ or when the latency of T2 targets is explicitly cued (Martens & Johnson, 2005; Nieuwenstein, Chun, van der Lubbe, & Hooge, 2005).

The second question that we addressed was related to the extent to which nonconscious stimuli can trigger prediction error responses, as measured with EEG. Over the last 20 years, we and others have shown that nonconscious information processing is rather sophisticated ^57,58^, and that a diverse range of high-level cognitive processes can unfold unconsciously, including motivational processing ^59–61^, semantic processing ^62,63^, decision making ^61,64^ and cognitive control ^65,66^. Interestingly, in Experiment 2 we found evidence that predictions that are violated by a nonconscious stimulus trigger a stronger negative fronto-central ERP component than predictions that are confirmed. This neural event was similar for trials on which T2 was seen and on trials where T2 was missed, highlighting that conscious awareness of a stimulus is not a prerequisite for it to trigger a prediction error response. This effect may reflect a mismatch signal, similar to the mismatch negativity ^29^, which is a negative deflection following oddball stimuli that develops 100-200 milliseconds after stimulus onset, resulting from violations of predictions. Sometimes this effect lasts longer, in some experiments until ~400 milliseconds, depending on the specifics of the task and stimulus material ^30,37,38^. While in terms of interpretation this effect is similar to a mismatch effect, its topography is slightly different than a typical visually evoked MMN, which generally peaks more posteriorly, although considerable variation in its topography has been reported ^30^. Alternatively, it is possible that the higher activation for valid compared to invalid trials corresponds to the frontal selection positivity (FSP), which is a well-known marker of non-spatial attentional processes ^67^. In our paradigm, this could be explained as improved attentional selection when predictions are confirmed. Although the exact nature of the observed component deserves future experimentation, the key finding is that the effect was independent of conscious access of T2 and purely depends on the validity of the prediction. This result is in line with studies that have shown context influences on nonconscious information processing ^68–71^, studies that have shown that the MMN can be observed when the prediction violations are unattended ^31,35,37,38,72^ and more generally evidence for relatively high-level processing of nonconscious stimuli ^58,73,74^. Nevertheless, the absence of interactions in the ERP is also somewhat surprising, because as noted earlier such interactions between prediction validity and conscious T2 detection were present in behavior. A neural basis for this effect should exist, but may be very subtle. Recently, a study by Aru et al. found early (<100 ms) differences in signal amplitude over posterior channels that predicted the behavioral benefit of prior knowledge on the detection of stimuli presented at the threshold of perception ^27^. Another potentially interesting signature to investigate could be the onset of components related to conscious perception ^22^ and how they relate to predictions. Moreover, it is possible that instead of signal strength, it is the signal-to-noise ratio or sharpness of the representation that is improved ^10^. Possibly, valid predictions do not just amplify or downregulate the neural response to a stimulus, but increase the specificity of the neural representation.

The final and third question regarded the necessity of a predictive stimulus itself to be consciously perceived to influence subsequent conscious access of stimuli in the attentional blink. In Experiment 3, we showed that conscious perception of T1 is a prerequisite for top-down predictive influences on conscious access to occur. On the subset of trials (~25%) where subjects did not see T1, there was no predictive influence of T1 on T2 detection performance. This result contrasts with findings from a recent study that suggested that some priors may operate nonconsciously (Chang, Baria, Flounders, & He, 2016). Chang and colleagues performed a study in which they presented participants with masked grey-scale natural scene images and found that the nonconscious processing of these images improved subsequent recognition of their degraded counterparts, so-called “Mooney images”, presented seconds later. One explanation for this difference is the fact that the priors on which the effects of Chang et al. relied are more automatic and hard-wired than the relatively arbitrary predictive relationships that people have to learn and actively use in our experiments. It is possible that lower-level, automatic predictions are more easily processed outside of awareness compared to the more active ones studied here.

Further, it is also possible that with more training on the predictive relationships in our task we would find nonconscious predictive effects. However, since subjects were already trained on the task on a separate day before performing the experimental session, this possibility seems unlikely. We did observe greater validity effects when subjects were made explicitly aware of the predictive nature of T1, suggesting that explicit knowledge of stimulus associations can facilitate the effects of stimulus-induced predictions. Finally, it should be noted that we did not test the full range of timing intervals between T1 and T2. It has been shown and proposed that the processing of nonconscious stimuli is relatively fleeting ^42,76^, so it is conceivable that the T1-T2 lags that we have used in our experiments may have been too long to observe predictive effects. However, recently it has been shown that the duration of nonconscious information processing may have been underestimated for a long time ^77^. Further, a significant attentional blink was observed on trials on which T1 was missed, indicating that attention was still captured by a missed T1 at the T1-T2 lags used in these experiments. This latter result is in line with evidence showing that nonconscious stimuli are able to trigger attentional capture ^78–80^ and with a study by Nieuwenstein and colleagues in which T2 detection was lower for T1’s that were missed compared to trials without T1 (in that experiment this effect was independent of lag (Nieuwenstein, Van der Burg, Theeuwes, Wyble, & Potter, 2009)).

In summary, three main conclusions can be drawn from the present series of studies. First, prediction confirmation, compared to violation, increases the likelihood of conscious awareness, suggesting that valid predictions amplify the perceptual strength of a stimulus. Second, nonconscious violations of conscious predictions are registered in the human brain, indicating relatively high-level processing of nonconscious stimuli. Third, however, predictions need to be implemented consciously to subsequently modulate conscious access. These results suggest a differential role of conscious awareness in the hierarchy of predictive processing, in which the active implementation of top-down predictions requires conscious awareness, whereas a conscious prediction and a nonconscious stimulus can interact to generate prediction errors. How these nonconscious prediction errors are used for updating future behavior and shaping trial-by-trial learning is a matter for future experimentation.

## Methods

### Participants

We tested 26 participants in Experiment 1 (20 females, age 19.5±1.3 years), 85 participants in Experiment 2 (49 females, age 21.9±3.0 years) and 34 participants in Experiment 3 (22 females, age 20.0±1.1 years). All participants were right-handed and had normal or corrected-to-normal vision.

For all experiments, participants for whom the minimum number of observations in one or more conditions was lower than 10, were excluded from analysis. Additionally, for Experiment 3 (EEG), we had to exclude 2 participants due to problems with the reference electrodes. In the end, this resulted in the inclusion of 25 participants for Experiment 1, 67 participants for Experiment 2 and 29 participants for Experiment 3.

The studies were approved by the local ethics committee of the University of Amsterdam and written informed consent was obtained from all participants according to the Declaration of Helsinki. Compensation was 20 Euros for Experiment 1, 25 euros for Experiment 2 and 30 Euros for Experiment 3, or equivalents in course credit.

### Materials

All stimuli were generated using the Psychophysics Toolbox ^82^ within a MATLAB environment (MathWorks, Natick, MA, USA). Stimuli were displayed on an ASUS LCD monitor (1920 x 1080 pixels, 120Hz) on a black (RGB: [0 0 0]) background while participants were seated in a dimly lit room, approximately 70 cm away from the screen.

### Procedure and stimuli

Participants performed an attentional blink task ^45^, in which on every trial a rapid series of visual stimuli was presented consisting of a sequence of 17 uppercase letters drawn from the alphabet but excluding the letters I, L, O, Q, U, and V. Letters were presented at fixation in a mono-spaced font (font size: 40) for 92 ms each.

### Experiment 1

Participants were instructed to detect target letters within the rapid serial visual presentation (RSVP). The first target (T1: G or H) was always presented at the fifth position of the RSVP. On most trials (80%) it was followed by a second target (T2: D or K) at lag 2, lag 4 or lag 10 (respectively 183, 367 or 917 ms later). Each lag was equally likely. T1 was presented in green (RGB: [0 255 0]), while T2 and the distractor letters were white (RGB: [255 255 255]).

Crucially, there was a predictive relationship between the two targets (Fig. 1A). Namely, when T2 was present, the identity of T1 (e.g. G) predicted which T2 was likely (75%, e.g. D) or unlikely (25%, e.g. K) to appear. On the 20% remaining trials without a T2 a random distractor letter was presented at the T2-timepoint. The mapping of T1 and T2 was counterbalanced over participants, so that for half of the participants the most likely target combinations were G-D and H-K while for the other half G-K and H-D were most likely. To be able to distinguish different lags in the absence of a T2 stimulus, 4 grey squares (RGB: [200 200 200]; size: .35°; centered at 1.30^°^ from fixation) were always presented around the stimulus (T2 or distractor) at the T2-timepoint. Participants were instructed to use the timing information this cue provided when making decisions about T2.

Following a 150 ms blank period at the end of the RSVP, participants gave their responses. First, they indicated whether or not they had seen any T2 by pressing the left or right shift key on the keyboard. The mapping between the keys and the response options was randomized per trial to decouple participants’ responses from the decision they had to make. Then they were asked to make a forced choice judgment about the T2 letter (D or K) that was presented by typing in this letter. Finally, they made a similar response about the identity of T1 (G or H). We used long response timeout durations of 5s and participants were instructed to value accuracy over response speed. The inter-trial interval, as defined by the time between the last response and the onset of the stream, was 500-750 ms.

The experiment consisted of two one-hour sessions on separate days within one week. In the first session, participants received instructions about the task and subsequently performed the task for 6 blocks of 75 trials (total 300 trials). The goal of the training session was to familiarize participants with the task. Besides, since we did not instruct participants about the predictive relationship between T1 and T2, some practice on the task was required for them to learn this relationship. In the second session, participants first received a summary of the instructions, after which the actual experiment started. Participants performed 6 blocks of 90 trials (total of 540 trials) of the AB task. The first three participants performed 6 blocks of 105 trials (630 trials). In both sessions participants received summary feedback about their performance at the end of each block, followed by a short break.

### Experiment 2 (EEG)

The task in the EEG experiment was the same as in Experiment 1, except that in Experiment 2, we only asked participant to give one response by typing in the target letters they observed. In addition, we only used two different lags: lag 3 (275 ms; 2/3rd of trials) and lag 10 (917 ms; 1/3rd of trials). To further increase the number of trials, the ITI range was reduced to 200-400 ms.

Again, the experiment consisted of two different sessions within one week. The first session (1 hour) consisted of instructions followed by extensive training (720 trials over 6 blocks) on the task. Participants were not explicitly informed about the predictive relationship between the targets. In the second session (2 hours) we first prepared the participant for the EEG measurements (see below) and gave brief instructions about the task. Then, participants performed 12 blocks of 120 trials (total 1440 trials) of the AB task.

### Experiment 3

To investigate the importance of T1 detection for prediction effects on conscious access, we adjusted the task we used in Experiment 1 to decrease the visibility of T1 (Fig. 3A). We now presented T1 in white instead of green, to make it stand out less among the other stimuli. Furthermore, T1 duration was staircased per participant such that participants could report T1 on roughly 75% of the trials. Starting in the second half of the training and continuing in the experimental session, after each block T1 duration was decreased by one frame (8 ms) if performance was higher than 85% and increased by one frame if performance was lower than 65%. To ensure T1 duration would not deviate too much from the duration of other stimuli, T1 duration was only allowed to be in the range of 42-142 ms (max. 50 ms different from other stimuli). The median duration of T1 in the second session was 125 ms. On 20% of trials no T1 was presented and a random distractor letter was presented instead. When both targets were present, T1 predicted which T2 was likely to follow with 75% accuracy.

We made a few changes to the task design to increase the efficiency of the design. The ITI was reduced to values between 300-500 ms. In addition, we only asked participants for one response. They were asked to type in any target letter they had seen during the trial and refrain from typing in a T1 and/or T2 letter when they did not see any. The response was confirmed by pressing the space bar on the keyboard or when a timeout of 4s had passed. To further increase the number of trials per condition, we decided to use only lag 3 (2/3rd of trials) and lag 10 (1/3rd of trials). Because T1 duration was staircased on an individual basis, the T1-T2 SOA differed between participants. On average, lag 3 corresponded to an SOA of 308 ms while lag 10 corresponded to an SOA of 950 ms.

Finally, we manipulated the instructions we gave to participants in order to see to what extent explicit knowledge of the relationship between T1 and T2 affected our results. As in Experiment 1, we tested participants during two separate sessions within one week. The first group of the participants (N=25) did not receive any explicit instruction about this relationship, similar to Experiment 1. The second group of participants (N=19) received explicit instructions about the T1-T2 relationship at the start of the second session, and a third group of participants (N=23) received those instructions already at the start of their first session.

The first session (1 hour) was used for instructions and training the participants on the task (10 x 75 trials). The experimental session in which participants performed the AB task lasted 1.5 hour and contained 16 blocks of 75 trials (1200 trials).

#### Behavioral analysis

Preparatory steps were done with in-house MATLAB scripts. Statistical analyses (repeated measures ANOVAs and paired t-tests) were performed using JASP software ^83^. In situations where a specifically tested hypothesis did not yield a significant result, we used a Bayesian equivalent of the same test to quantify the evidence for the null-hypothesis ^84,85^. In those cases, using JASP’s default Cauchy prior, Bayes Factors (BF) were computed for each effect. To increase the interpretability in analyses with multiple factors, we used Bayesian model averaging to get a single BF for each effect in ANOVA’s. This BF is the change from prior to posterior inclusion odds, and can intuitively be understood as the amount of evidence the data gives for including an experimental factor in a model of the data. The BF will either converge to zero when the factor should not be included, or to infinity when it should be included in the model. Values close to one indicate that there is not enough evidence for either conclusion. We use the conventions from Jeffreys (1967) to interpret the effect sizes of our Bayesian analyses.

### Experiment 1

In our behavioral analyses we looked at the T2 detection performance, given that T1 was correctly identified. A detection response was considered to be correct when (1) the participant indicated no T2 was present when no T2 was presented or (2) the participant correctly indicated a T2 was present and subsequently reported the correct target letter. Trials where one of the responses was missing were deleted from all analyses. Percentage correct was used in a 3 x 2 repeated measures ANOVA with the factors lag (lag 2, lag 4, lag 10) and prediction (valid, invalid). In a control analysis, we repeated our analyses for Experiment 1 based on the T2 detection responses (ignoring the accuracy of the T2 identification) as dependent variable (see also Results). Since the seen/miss response is orthogonal to the specific predictions about target letters, this analysis rules out simple response biases as a potential cause of our effects.

### Experiment 2 (EEG)

The behavioral analyses for the EEG experiment were similar to those for Experiment 1. However, the factor lag had only 2 levels (lag 3, lag 10). Percentage correct T2 detection was computed as in Experiment 1 using only the trials on which T1 was correctly reported. A response was considered to be correct when the letter a participant entered was the letter that was presented or when a participant refrained from entering a letter when none was presented.

### Experiment 3

In this experiment, participants gave only one response by typing in the target they had perceived. Trials on which no response was given or on which an impossible response was given (e.g. two T1 targets reported) were excluded from analyses. For T1 and T2 separately, we assessed the accuracy of the responses. The definition of correct and incorrect responses was the same as in Experiment 2.

Subsequently, T2 percentage correct detection was used in a 2 x 2 x 2 x 3 mixed ANOVA with the within-subject factors lag (lag 3, lag 10), prediction (valid, invalid) and T1 visibility (T1 seen, T1 missed) and the between-subject factor instruction (none, start session 2, start session 1). As mentioned before, this between-subject factor was included to find out whether predictive effects would be modulated by explicit knowledge of the relation between T1 and T2. To investigate the effect of T1 visibility in more detail, we followed up the main analyses by other mixed ANOVAs in which we first split up the dataset based on T1 visibility. In situations where we found interactions with the factor instruction, we compared the effects of lag and prediction separately per instruction condition using repeated measures ANOVAs and paired-samples t-tests.

Finally, to test for an interaction between prediction validity and lag, we combined behavioral data from all experiments in a post-hoc analysis. Only trials on which T1 was correctly identified were used. For Experiment 1 we averaged data for lag 2 and lag 4 to create an average “short lag” condition. Subsequently, these data were entered into a 2 x 2 x 3 mixed ANOVA with the within-subject factors lag (short, long) and prediction (valid, invalid) and the between-subject factor experiment (Experiment 1, Experiment 2, Experiment 3).

#### Electroencephalographic measurements

EEG was recorded with a BioSemi ActiveTwo system and sampled at 512 Hz (BioSemi, Amsterdam, The Netherlands). Sixty-four scalp electrodes were measured, along with two reference electrodes on the earlobes and four electrodes measuring horizontal and vertical eye movements. After data acquisition, EEG data was pre-processed with the FieldTrip toolbox for MATLAB ^87^. First, data were re-referenced to the linked earlobes, high-pass filtered at 0.01 Hz, and epoched from -0.750 to 1s surrounding the onset of T2. Data were visually inspected and trials and/or channels containing artifacts not related to eye blinks were manually removed, resulting in deletion of on average 9.1% (±3.9%) of trials and 2.0 (±1.7) channels. Independent component analysis was used to identify components related to eye blinks or other artifacts that could easily be distinguished from other EEG signals. After the independent component analysis, previously deleted channels were reconstructed based on a nearest neighbor approach. Trials were baseline corrected to the average amplitude prior to T1 onset (-0.750 to -0.275). As a final step, we applied a 40Hz low-pass filter to the trial data, after which ERPs were created separately for each condition of interest.

#### Electroencephalographic analyses

All EEG analyses are based exclusively on trials where T2 appeared at lag 3 and T1 was correctly identified. We used a combination of Fieldtrip ^87^ and in-house MATLAB scripts to perform our analyses. As a first step, we performed cluster-based permutation tests ^49^ on the time-window 0-750 ms from stimulus onset to isolate significant ERP events relating to prediction validity (irrespective of T2 visibility) or T2 visibility (irrespective of validity) or the interaction between those factors. Next, we used an in-house build MATLAB script to isolate the significant events as clusters in time and space. For this purpose, we computed an average difference wave over all channels that were part of the cluster at any point in time. Subsequently, the onset and offset of a cluster were defined as the time period around the maximum difference where the difference did not drop below 50% of this maximum and where at least one channel showed a significant effect. We then selected the 10 channels that showed the largest effect in this time-window. One of the observed events reflected a mixture of the traditionally observed P3a and P3b components ^50,88^. Therefore, we split the event into two clusters by manually selecting either the 32 most anterior or 32 most posterior EEG channels (from the central midline) before running the cluster selection procedure.

As an alternative way to establish potential interactions between T2 detection and validity, we inspected the clusters that were isolated in the previous step in more detail. This may be a more powerful (but also less sensitive) way to detect small effects, because data is averaged over time-points and channels. Within each of the clusters, we performed a 2 x 2 repeated measures ANOVA (and its Bayesian equivalent, see also Behavioral Analysis) with the factors T2 detection (seen, missed) and prediction validity (valid, invalid) on the cluster data averaged over channels and time. To prevent double dipping, in each analysis we only considered the effects orthogonal to the one that was used to define the cluster (e.g. not testing the effect of prediction in a cluster defined based on the prediction effect).

## Acknowledgements

This research was supported by the Netherlands Organization for Scientific Research (NWO VENI (451-11-007) awarded to SvG; NWO VIDI (452-13-016) awarded to FdL), the European Research Council (ERC-2015-STG_679399 awarded to HAS) and the James S McDonnell Foundation (Understanding Human Cognition, 220020373, awarded to FdL). We thank Doris Dijksterhuis, Sjoerd Manger and Thomas Dolman for their valuable assistance with data acquisition. We thank Timo Stein and Josipa Alilovic for valuable comments on a previous draft of this manuscript.

## Supplementary Results

### Experiment 2: Visibility effects

Here we report the ERP events that are associated with T2 visibility (irrespective of prediction validity) and the interaction between prediction and visibility. To analyze the events related to T2 visibility, we examined the difference in ERPs following seen and missed T2s using a cluster-based permutation test (see Sup. Fig. 1), thereby correcting for multiple comparisons across both time and (electrode) space ^49^. Similar to previous findings (e.g. Sergent et al., 2005), this revealed two significant events. First, a significant negative difference could be observed over (left) posterior electrodes from 170-355 ms after T2 onset (p=0.010; Sup. Fig. 1A). This event was followed by a significant long-lasting positive event (p<0.001), reflecting a mixture of the P3a and P3b components, extending frontal and central electrodes.

Subsequently, we wanted to have a closer look at the interactions between conscious access and prediction validity. Therefore, we analyzed the ERP events that were isolated in the previous step in more detail (Sup. Fig. 1B-G). For this analysis we isolated the traditionally observed AB-related P3a and P3b ERP components from the long-lasting positive ERP event that differentiated between seen and missed T2s ^50^. Doing so resulted in an early positive P3a cluster (Sup. Fig. 1D) over fronto-central channels that was significant between 395-586 ms and a somewhat later positive P3b cluster (Sup. Fig. 1F) over more posterior parietal channels, which was significant between 445-611 ms.

Next, we performed repeated measures ANOVAs in each of these clusters. For none of the events we found evidence of such interactions (early left-posterior event: F_1,28_=0.29, p=0.597, BF=.260; P3a: F_1,28_=1.56, p=0.222, BF=0.230; P3b: F_1,28_=2.10, p=0.159, BF=0.296). This suggests that for the events related to T2 detection the amplitude of the signal did not depend on the validity of the predictions, though the BF values suggest that the evidence for the absence of such interactions is moderate at best.

In sum, similar to earlier studies we found a sequence of events differentiating trials on which T2 was seen from trials on which it was not perceived ^50^. First, seen T2s evoked a stronger posterior negativity, which was followed by a large central positive effect reflecting a mixture of the P3a and P3b components. Especially these later positive components have previously been related to conscious access ^50,89^ and metacognition ^90^, though recent investigations show it may also reflect even later more cognitive processes, merely arising as a consequence of becoming consciously aware of information ^73,91^. We did not find evidence that the amplitude of these ERP events was modulated by prediction validity. This may suggest that once a stimulus was perceived consciously, it is irrelevant whether or not the prediction was valid.

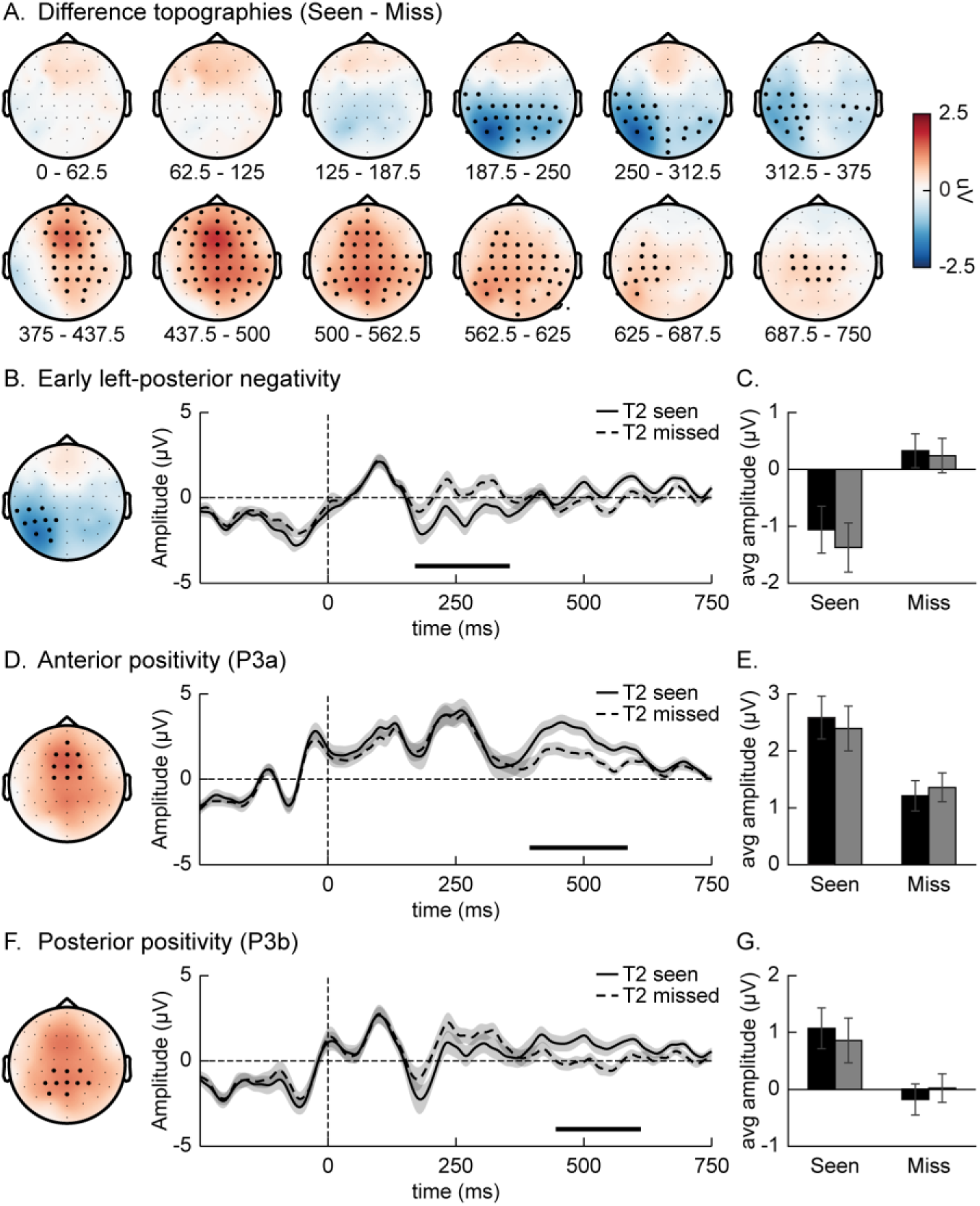
Supplementary Figure S1. ERP effects related to T2 visibility analyses. **(A)** Topographic maps showing the difference between seen and missed T2s over time (0 = T2 onset). Cluster-based permutation tests were used to isolate the significant events while correcting for multiple comparisons across time and (electrode) space. On each head map, channels showing a significant difference for at least 50% of its time window are highlighted. Three events were isolated based on the permutation tests. **(B,D,F)** For each of the events individually, the average ERP time-course of the 10 channels shown on the headmap on the left, seperately for T2 seen and T2 missed conditions is shown. The significant time-window is marked by a black line above the x-axis. **(C,E,G)** Bar graphs showing the average amplitude of the four conditions (visibility x prediciton) for the significant neural events shown in **B,D,F**. In all plots error bars represent SEM.

